# Low-cost, scalable, and automated fluid sampling for fluidics applications

**DOI:** 10.1101/2021.01.27.428538

**Authors:** A. Sina Booeshaghi, Yeokyoung (Anne) Kil, Kyung Hoi (Joseph) Min, Jase Gehring, Lior Pachter

## Abstract

We present colosseum, a low-cost, modular, and automated fluid sampling device for scalable fluidic applications. The colosseum fraction collector uses a single motor, can be built for less than $100 using off-the-shelf and 3D-printed components, and can be assembled in less than an hour. Build Instructions and source files are available at https://github.com/pachterlab/colosseum.

## Introduction

Fraction collectors that sample from a microfluidic stream (Blume et al., 2015), are preferable to manual collection that can be tedious and introduce human error (Jessop-Fabre and Sonnenschein, 2019). Commonly used in fast protein liquid chromatography (FPLC), typical fraction collectors consist of a rotating rack loaded with containers and a distributing arm for collecting fixed volumes of fluid (Madadlou et al., 2017; Polson, 1961). Most laboratories currently rely on commercial fraction collectors, which are expensive and difficult to customize (Supplementary Table 1). To reduce cost and facilitate custom applications, a number of open-source fraction collectors have been developed, e.g. (Caputo et al., 2020; Longwell and Fordyce, 2020). These devices, while less expensive, continue to rely on complex engineering designs and parts that may be difficult to source and manufacture, thus driving costs higher, lengthening the assembly process, and complicating operation.

We have designed and built a simple, low-cost, and modular fraction collector that is easy to assemble and use. This open-source fraction collector, which we call colosseum, is based on design principles for modular, robust, open-source hardware (Booeshaghi et al., 2019), and offers advantages to commercial systems by virtue of being significantly less expensive and easily customizable.

The colosseum fraction collector can be assembled in less than an hour and costs $67.02. Unlike the micrIO (Longwell and Fordyce, 2020), which is built from parts of a salvaged Illumina Genome Analyzer that costs $1500, the colosseum fraction collector uses off-the-shelf and 3D-printed parts (Supplementary Table 2). The LEGO MINDSTORM fraction collector (Caputo et al., 2020) costs $500, and while it uses more commonly available components, it still requires cutting and bending of steel C-channel. Furthermore, most fraction collectors require the use of multiple axes to position a dispenser head over a reservoir. Control of such a system can require communicating with and driving up to three separate motors in tandem. The colosseum fraction collector is based on a simpler design where a mechanical coupling between the motor, the tube rack, and the dispenser arm enables rotation of the rack and position of the arm with only one motor. Designing around a single motor simplifies operation, and reduces cost, complexity, and assembly time.

## Results

The colosseum fraction collector consists of four 3D-printed components, two rotary shafts, five rubber feet, one stepper motor, an Arduino, and a motor controller (Figure 1a,b). We chose the spiral tube layout (Figure 1c) instead of the rectangular tube layout of previously published fraction collectors as it enables serial fraction collection with only one motor. By coupling the dispenser arm to the tube rack with a slot-cam mechanical coupling (Figure 1d) we constrained the rotation of the tube rack and movement of the dispenser arm to rotation of a single stepper motor located in the base of the fraction collector (Figure 1e).

**Figure 1:**
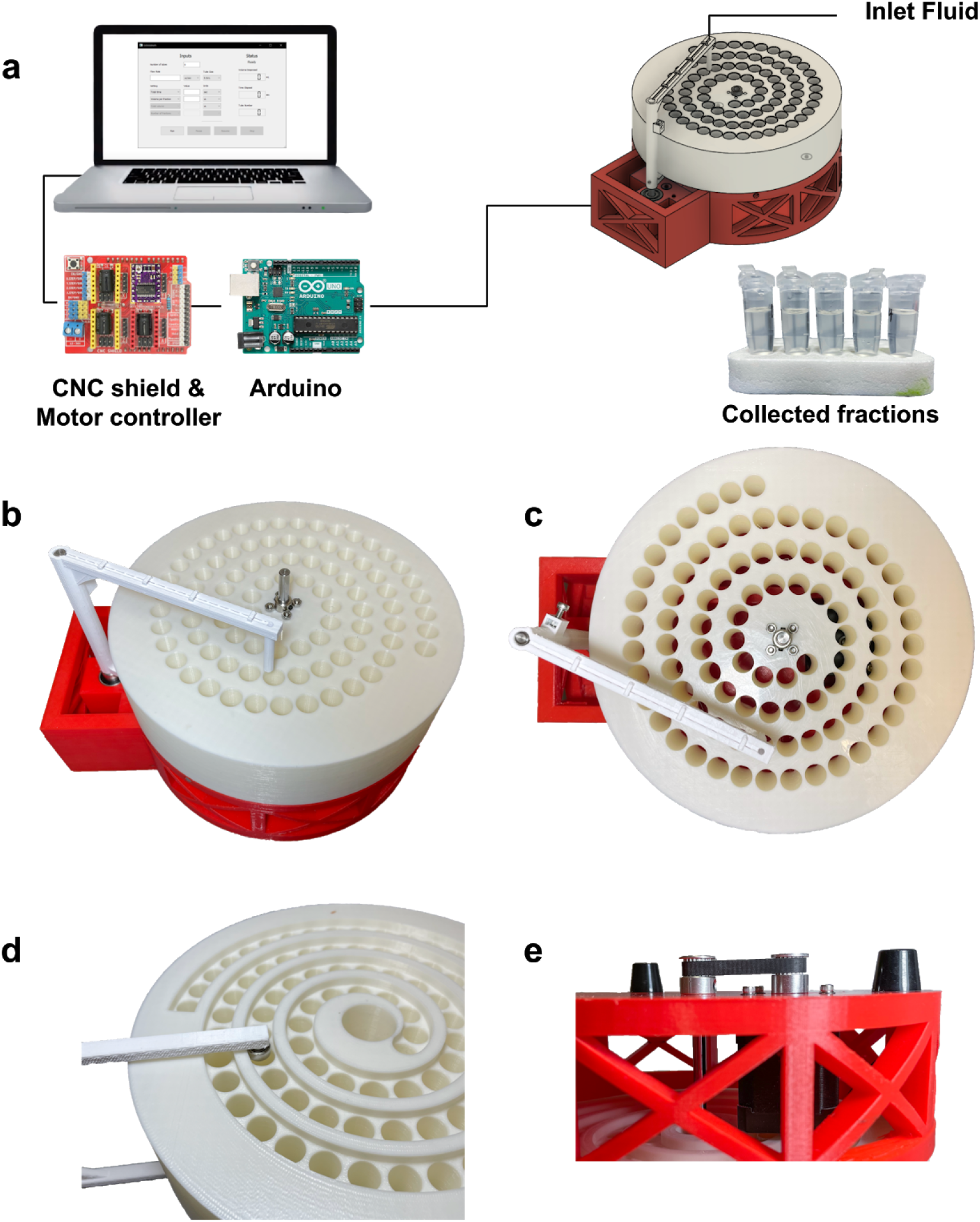
(a) The colosseum fraction collector (left) is controlled by a single motor. A motor controller shield (red) is connected to an Arduino Uno (blue) and drives the motor. The computer’s Graphical User Interface (right) and Python backend sends motor movement instructions to the Arduino. The Arduino-motor controller then sends those instructions to the motor. A motor located in the base turns the shaft of the tube rack. Grooves in the bottom of the fraction collector constrain the dispenser arm to rotate in tandem. (b) Angled view, (c) top view, (d) bottom view of mechanical coupling between the dispenser arm and tube rack, (e) side view of mechanical coupling of motor and tube rack.

The device is modular: each component can be developed, tested, and fabricated separately using mutually compatible interfaces. The tube rack fits 1.5 mL Eppendorf tubes and can easily be modified to accept tubes of varying sizes under the constraint that they follow the spiral pattern (Figure 2a). The tube rack fits 88 tubes with a packing efficiency of 60.2% relative to the optimal packing of 146 tubes on a circular disk the same size as the area available for placing tubes on the tube rack (Supplementary Figure 1; Specht, 2018). In addition, the dispenser arm can be modified to accept connectors and tubing of various sizes to enable parallel dispensing.

**Figure 2:**
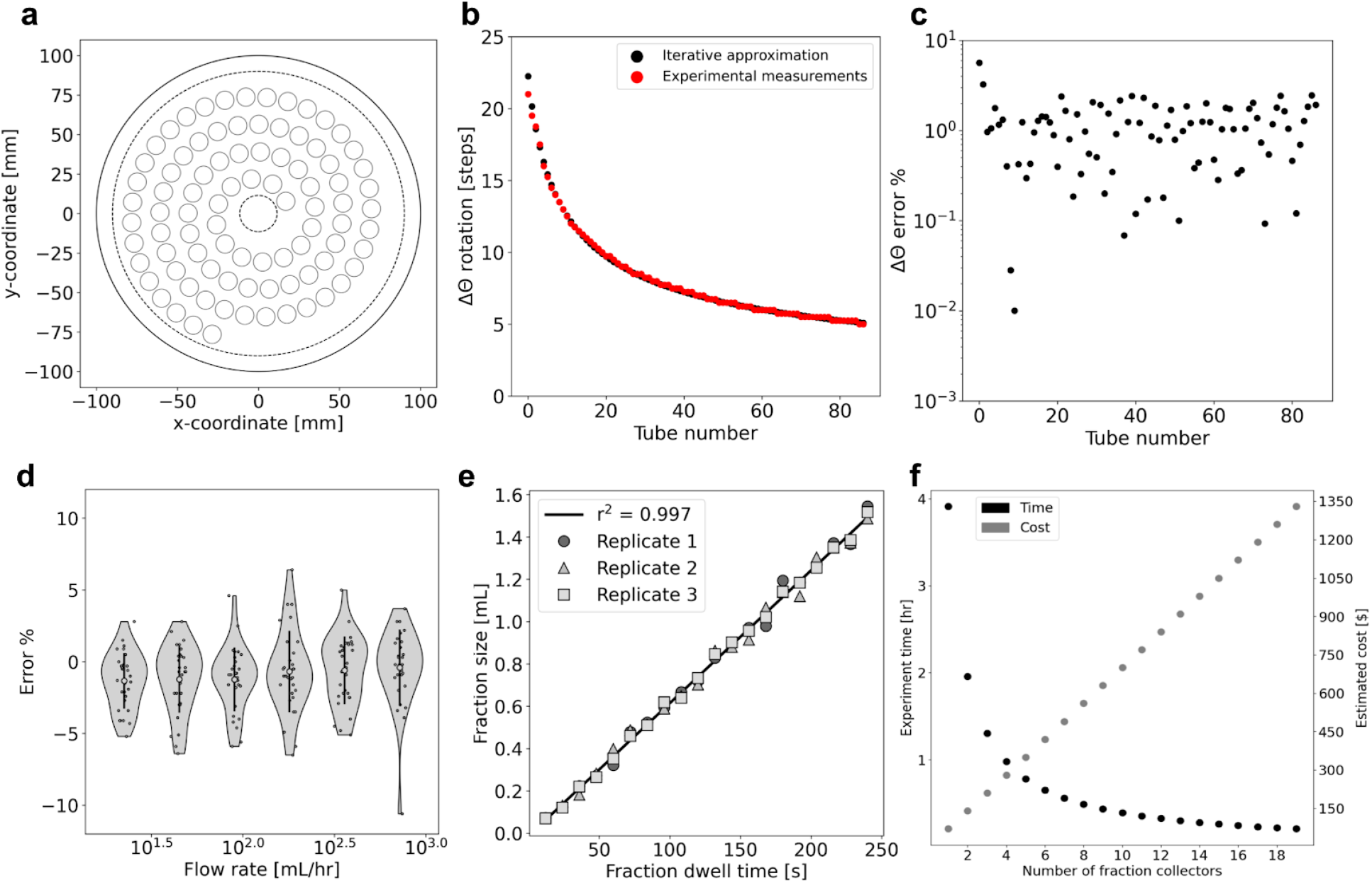
(a) Tube placement on the colosseum is defined by an Archimedean spiral with tubes distributed 13 mm apart along the spiral and with 17.39 mm distance between subsequent arms of the spiral. The dotted-line innermost circle corresponds to the area on the tube rack designated for the set screws on the center shaft. The larger dotted-line circle corresponds to the area available for tubes. The solid outer circle corresponds to the tube rack boundary. (b) The tubes are placed uniformly along the spiral where the arc length between any two tubes is constant, but the rotational displacement between any two tubes is nonconstant. (c) Iterative approximation to the tube locations is similar to the measured tube locations. (d) The error in the fraction size for 88 samples across a range of flow rates. (e) The fraction size increases with increasing dwell time for a constant flow rate and the Spearman correlation of the means is 0.997. (f) Multiple fraction collectors enable parallel collection which drastically decreases experimental time at a marginal increase in cost. [[Code a,b,c](https://github.com/pachterlab/BKMGP_2021/blob/main/analysis/archimedian_spiral.ipynb), [Code d](https://github.com/pachterlab/BKMGP_2021/blob/main/analysis/varying_flowrate.ipynb), [Code e](https://github.com/pachterlab/BKMGP_2021/blob/main/analysis/constant_flowrate.ipynb), [Code f](https://github.com/pachterlab/BKMGP_2021/blob/main/analysis/scalability.ipynb)]

The device is controlled by a graphical user interface (GUI) that communicates with an Arduino, CNC motor shield, stepper motor driver, and software adapted from the poseidon syringe pump (Supplementary Figure 2; Booeshaghi et al., 2019). Experiment parameters such as flow rate, total volume, total time, volume per fraction, and number of fractions are input by the user in the GUI (Supplementary Table 3) and the python back-end structures and sends Arduino-interpretable commands to the Arduino for execution. The GUI can be installed with the pip package-management tool and run with a single command on Mac, Linux, or Windows.

To ensure that commands set by the GUI correctly align the dispenser arm with each collection tube, we measured and converted the angle between pairs of tubes to motor steps, and programmed this list of angular displacements into the control software (Methods). We also used a simple iterative scheme to approximate the position of equally spaced points along an Archimedean spiral and compared it to our measurements. We found high concordance in the angular displacements (Figure 2b,c). This allows us to programmatically generate arbitrary spiral motor displacements based on the distances between successive tubes and distances between successive arms of the spiral.

In order to characterize collection errors across a range of flow rates commonly used in microfluidics and FPLC (Cheri et al., 2014; Westman et al., 1987), we sampled 180 fractions over six flow rates ranging from 22.5 mL/hr to 720 mL/hr. We found that the collection errors were within ±6.5%, with one sample having −10.6% error due it being the first fraction collected. These data suggest that the use of the colosseum system with the poseidon syringe pump results in accurately collected fractions (Figure 2d). Next, we sought to assess the fraction collecting performance over an increasing amount of sample volume, as is commonly performed in gradient elution series (Polson, 1961). For a fixed flow rate, we collected 20 fractions with 12-second increments in collection time per tube over the course of 42 minutes in three replicates (Figure 2e). We found that the collected fractions closely followed the expected fraction amount with a Spearman correlation of 0.997, showing that the colosseum fraction collector can be used to accurately collect gradient elution series.

## Discussion

We have demonstrated a low-cost, modular, and automated fraction collector that uses 3D-printed parts and off-the-shelf components, can be built in an hour, and is simple to run. We show how colosseum samples fluid accurately over a wide-range of flow rates making it useful for microfluidics experiments and FPLC. The low cost of our device could enable several instruments to run in parallel. For example a single control board can in principle run multiple fraction collectors and syringe pumps thus facilitating large-scale experiments (Figure 2f). We have also thoroughly documented the build process with instructional README’s and videos (Supplementary Figure 3), and we have made all of the results described in this paper reproducible on Google Colab.

## Methods

We designed the colosseum fraction collector by following basic principles of open-source hardware design (Booeshaghi et al., 2019).

### Part design

The fraction collector consists of four 3D-printed parts: a base, base plate, dispenser arm, and tube rack. The base holds the base plate, dispenser arm, and tube rack in place with additional hardware. The base plate acts as a horizontal support for the main rotary shaft, with rotational bearings that support the shaft in two places. The dispenser arm consists of two connected parts: the top part of the arm holds the fluid tubing and the bottom part acts as a cam follower that follows the spiral track on the bottom of the tube rack. Collection tubes are placed in the tube rack and are organized in a spiral pattern that mirrors the pattern the dispenser arm follows during rotation. The tube rack is constrained to the shaft with a flange coupling set screw and is mechanically coupled to the motor with a timing belt so that rotation of the motor results in rotation of the tube rack and the dispenser arm.

After numerous sketch iterations, we used Fusion 360 (Song et al., 2018) to generate a 3D model of the device and added dimensional tolerances of +3-5% to all parts to account for variance in 3D printing.

### Part fabrication

STL files were generated using the 3D part models from Fusion 360. To prepare the appropriate files for 3D printing, Simplify3D (Simplify3D) was used to slice the STL model and generate GCode with 10% infill and 0.2mm layer height. Parts were printed on a Prusa i3 Mk3 3D printer (Prusa Research). GCode was loaded onto an SD card and the parts were printed using 1.75mm diameter PLA filament with a nozzle temperature of 215°C and a bed temperature of 60°C. All parts were printed at 10% infill. STL files for all parts can be found in the [GitHub repository](https://github.com/pachterlab/colosseum/tree/master/HARDWARE). The time to print all parts separately was approximately 73 hours, but may vary depending on the printer model used and the print settings (Supplementary Table 2). All parts required to assemble the colosseum fraction collector can be found in the [bill of materials](https://github.com/pachterlab/BKMGP_2021/blob/main/HARDWARE/BOM.xlsx).

### Device assembly

A complete guide on how to assemble the colosseum fraction collector can be found on [YouTube](https://youtu.be/yG7ECh5GO0o) (Supplementary Figure 3).

The assembly of the device starts with the base. Five rubber feet are screwed onto the bottom of the base to stabilize the device and to ensure that the timing pulleys on the motor and the tube rack shaft are elevated and free of obstruction. A timing belt pulley is secured to the shaft of a Nema 17 motor and the motor is then screwed onto the floor of the base. The tube-rack shaft is also inserted into the floor of the base along with a bearing that acts to stabilize the shaft. A timing belt pulley is secured to the shaft and couples its rotation with that of the motor. The motor and the shaft are connected by a timing belt of length 120 mm. The mounting holes in the base for the motor are designed so that the user can adjust the distance between the two timing pulleys in order to prevent slippage of the timing belt. Additionally, washers are inserted in between the base floor and the screws holding the motors so that the plastic of the base does not get worn out over time.The base plate is then screwed onto the floor of the base using M5 screws and nuts.

The dispenser arm, which is secured to a shaft with an M3 set screw, is placed into the base plate along with a bearing. A torsion spring is placed on the shaft, between the dispenser arm and the base plate to lessen slack between the dispenser cam follower and the tube rack spiral groove. The tube rack shaft is then inserted into the tube rack and secured in place with a flange coupling set screw.

The motor cables are routed through the side of the base and connected to the Arduino. The Arduino is connected to a CNC shield and DRV8825 Pololu motor controller. The Arduino is also connected to a computer. This allows the user to send and receive signals to the motor via serial commands. The Arduino receives 12V DC power from a power supply.

### User Interface

The graphical user interface (GUI) translates the parameters set by the user into motor commands sent to the Arduino. The Arduino runs our custom firmware, pegasus https://github.com/pachterlab/pegasus, which sends command strings to the motor controller which in turn sends pulse-width-modulated signals to the motor (Supplementary Figure 2). The GUI is written in Python using Qt, an open-source, cross-platform GUI framework. All packages related to the GUI are pip-installable and the GUI can be launched with a single command. The GUI consists of two parts: parameter inputs and a status monitor, the latter of which displays the total volume dispensed, time elapsed, and current tube location. Upon opening the GUI, users are prompted to connect to an Arduino. To run the colosseum fraction collector, users must specify three parameters: the flow rate, total time or total volume, and volume per fraction or number of fractions (Supplementary Table 3). The remaining parameters are calculated using the ones provided. In addition to these parameters, users must also specify the tube size to ensure that the fraction size will not be greater than the capacity of the tube. Users can operate colosseum by pressing the run, pause, resume, and stop buttons in the GUI. All software required to run the colosseum fraction collector is freely available on Github under an open source BSD-2-Clause License.

Python 3.6 code is used on the back end to interpret user input from the GUI and send custom commands to the Arduino, accordingly. Parameters from the GUI are translated into dwell time per tube and number of tubes to fill. The angle between each tube in the spiral was measured on Fusion 360 using the Inspect tool, saved as a csv file (Supplementary Table 4), and specified in the Python backend. These angles are then converted into the number of steps the motor must rotate. The motor stops rotating at each tube location for a specified amount of time in order to dispense the fluid into the tube. The motor then moves for a set number of steps to reach the next tube. The status monitor displays the amount of total volume dispensed, how much time has elapsed since the start of the experiment and which tube the fraction is being dispensed into.

### Testing and Validation

We tested the functionality of the device with numerous experiments where tap water is flown in at a set flow rate, or varying flow rate. We used the poseidon syringe pump, a 60 mL syringe, microfluidic tygon tubing and 1.5 mL Eppendorf tubes to pump fluid to the colosseum. The poseidon syringe pump was controlled with the pegasus software. For a varying number of flow rates and a set dwell time per tube for each flow rate (Supplementary Table 5), we collected 30 fractions and compared the fraction sizes to the predicted fraction size of 1 mL by weighing each tube before and after collection (Figure 2d). We used a 200x 1 mg analytical scale manufactured by Yae First Trading Co., ltd part number TEK-AB-0392 to measure the amount of collected fluid. In order to properly fit the Eppendorf tubes on the tube rack, we cut off the caps from the Eppendorf tubes before collecting fractions in them and put them back on for the final measurement making sure that the cap corresponded to the tube from which it was removed.

In follow up experiments we fixed the flow rate and linearly increased the collection time. For a fixed flow rate of 22.5 mL/hr and 20 fractions with 12-second increments in collection time per tube, we collected fractions and compared the observed fraction sizes to the predicted fraction sizes (Figure 2e). We used pegasus to run the colosseum with varying dwell times per tube. We estimated the cost and time for using k fraction collectors to show that these devices, when used in parallel, can reduce the experimentation time. For example, if we collect n fractions on each of k fraction collectors with a volume per fraction v and a constant flow rate f per collector then the time it takes to run this collection is t = n/k*v/f.

To test the accuracy of the measured angles between two successive tubes we used an iterative scheme to estimate the radius and angular position based of the polar form of Archimedean spiral of r=b*θ for a constant b. The radius and the arc length are used to update the angular position and then the angular position is used to update the radius.

Optimal packing was calculated with the “best known packings of equal circles in a circle” online tool (Specht 2018) with the outermost disk corresponding to the diameter of the area available for tube placement and the packing disks corresponding to the distance between tubes along the arc.

### Data analysis

All data analysis was performed with Python 3.7. Jupyter notebooks that run in Google Colab and all experimental data to reproduce Figure 2 can be found on our GitHub repository https://github.com/pachterlab/BKMGP_2021.

## Acknowledgments

We thank Justin Bois for naming the colosseum instrument. We also thank the Caltech Library Techlab for helping us 3D print parts. We thank Taleen Dilanyan for wet lab training and support. We thank Eduardo da Veiga Beltrame for assistance with 3D printing.

## Author contributions

ASB, YK, and LP designed the fraction collector. JG helped set instrument specifications. YK assembled and built the fraction collector and performed the experiments. ASB, YK, and KHM designed the GUI. KHM coded the GUI. ASB, YK, and KHM wrote the documentation. ASB and YK analyzed the data and made figures. ASB, YK, and LP wrote the manuscript.

## Conflicts of interest

The authors declare no conflicts of interest.

## Data & software availability

All data and software can be found in the GitHub repository: https://github.com/pachterlab/colosseum.

## Notes

https://github.com/pachterlab/colosseum

### Competing Interest Statement

The authors have declared no competing interest.

### Summary of Updates

Notations for 3D printing have been made consistent. Modified Figure 1 to include picture of collected fractions. Fixed typo "FLPC" in Discussion section. Updated BOM to include links to Aliexpress.

