## Supplementary Figures & Tables for "Low-cost, scalable, and automated fluid sampling for fluidics applications"

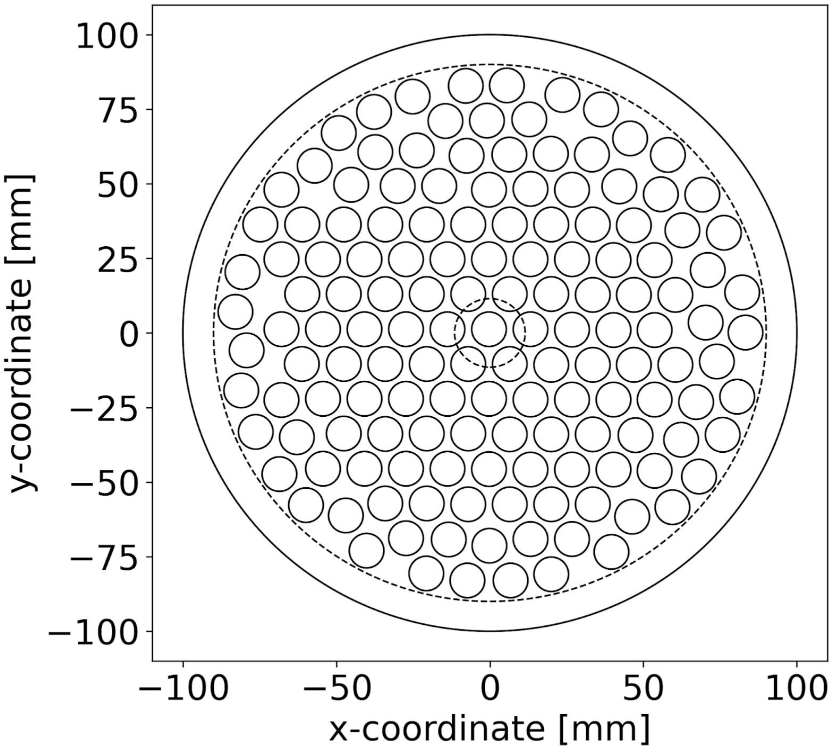


**Supplementary Figure 1:** Optimal packing of disks of diameter 13.5 mm (11 mm tube hole size plus 2.5 margin) on a disk of diameter 180 mm. The solid line corresponds to the outer diameter of the tube rack, the smaller dashed line corresponds to the effective area available for placing tubes, and the smallest dashed line corresponds to the empty area on colosseum where no tubes can be placed. [Code]

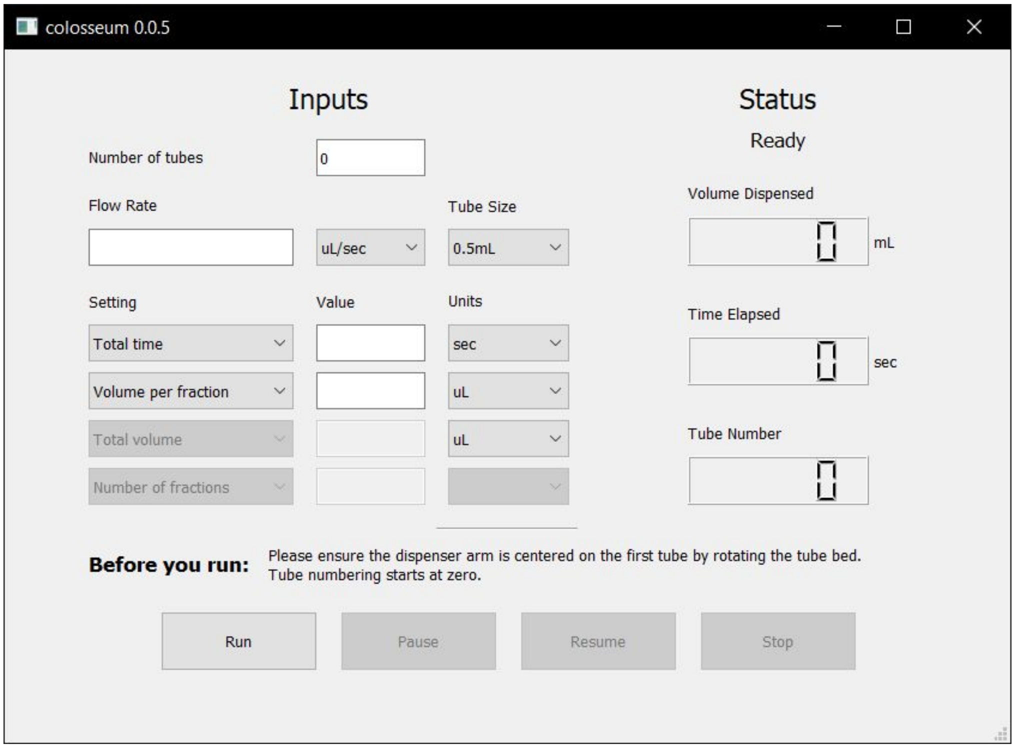


**Supplementary Figure 2:** The graphical user interface (GUI) of colosseum. The left panel displays input boxes for flow rate and collection parameters and the right panel displays experiment progress.

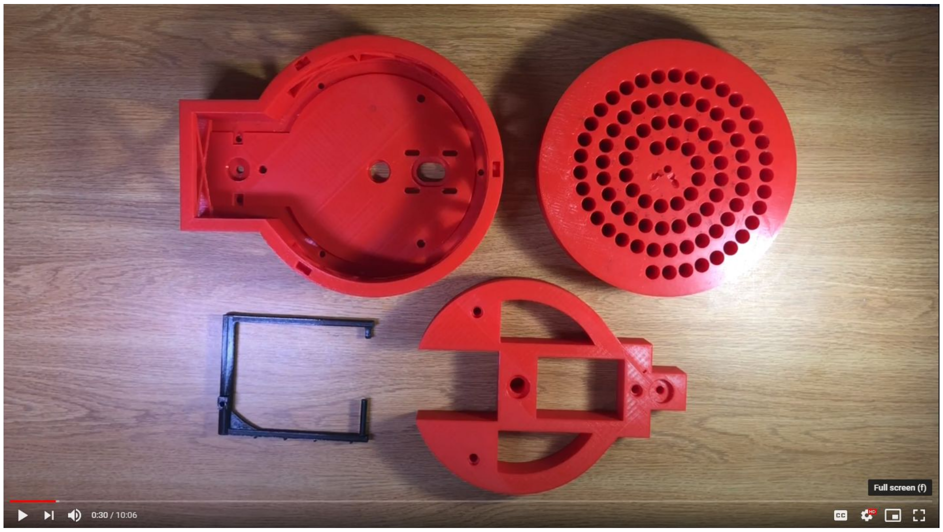


**Supplementary Figure 3:** Assembly video of colosseum. This video guides the user step-by-step through the entire assembly process**.** The video is linked to in the GitHub repository https://github.com/pachterlab/colosseum.

| Model | Capacity (# tubes) | Price (USD) |
|---|---|---|
| Cytiva Frac30 (1) | 30 | 1,615.00 |
| Eldex UFC (2) | 135 or 160 | 3,971.80 |
| Spectrum Spectra FC (3) | 174 | 3,393.00 |
| Buchi C-660 (4) | 12, 30, or 60 | 13,630.11 |
| Open-source | Customizable | <100 |

**Supplementary Table 1:** Costs and capacity of commercial fraction collectors. The costs are based on new, unused models. The capacity of each fraction collector is given by how many tubes the device can hold.

| Part name | Filament weight [length] | Print time | Supports |
|---|---|---|---|
| Tube Rack | 433.80 g [144.281 m] | 31 h 16 min | N |
| Dispenser Arm | 18.53 g [6.162 m] | 1 h 34 min | Y |
| Base | 271.45 g [90.285 m] | 19 h 4 min | Y |
| Base Plate | 174.70 g [58.105 m] | 11 h 26 min | N |
| Total | 898.48 g [298.833 m] | 73 h 30 min | |

**Supplementary Table 2:** Parts that require 3D printing, including, for each part, the amount of filament (weight and length) required to print, the print time, and whether support is required.

| | | | |
|---|---|---|---|
| Parameter 1 | Flow rate | | |
| Parameter 2 | Total time | OR | Total volume |
| Parameter 3 | Volume per fraction | OR | Number of fractions |

**Supplementary Table 3:** Table of input parameters for the GUI. The user must input three parameters: flow rate; total time or total volume; and volume per fraction or number of fractions. We limit users to three parameters to avoid overconstraining the system with conflicting parameters.

| Tube # | # of 1/4 steps |
|---|---|
| 0 | 84 |
| 1 | 78 |
| 2 | 75 |
| 3 | 70 |
| 4 | 64 |
| ... | ... |

**Supplementary Table 4:** The first five rows of the angles between each tube in the tube rack. The angular distances are reported as quarter-steps of the stepper motor. [Data]

| Flow rate (mL/hr) | Dwell time (s) |
|---|---|
| 720 | 5 |
| 360 | 10 |
| 180 | 20 |
| 90 | 40 |
| 45 | 80 |
| 22.5 | 160 |

**Supplementary Table 5:** Dwell time for each flow rate. To keep the expected fraction volume at 1 mL the flow rate is halved when the dwell time is doubled.
